# Loud music and the specific sound stress open the blood-brain barrier: new fundamental, biomedical, and social aspects

**DOI:** 10.1101/509042

**Authors:** O. Semyachkina-Glushkovskaya, V. Chekhonin, D. Bragin, O. Bragina, E. Vodovozova, A. Alekseeva, V. Salmin, A. Morgun, N. Malinovskaya, E. Osipova, E. Boytsova, A. Tohidpour, A. Shirokov, N. Navolokin, Y. Yang, C. Zhang, W. Feng, A. Abdurashitov, M. Ulanova, N. Shushunova, A. Khorovodov, A. Terskov, A. Esmat Shareef, A. Pavlov, Q. Luo, D. Zhu, V. Tuchin, J. Kurths.

## Abstract

The blood-brain barrier (BBB) poses a significant challenge for drug brain delivery. The limitation of our knowledge about the nature of BBB explains the slow progress in the therapy of brain diseases and absence of methods for drug brain delivery in the clinical practice.

Here we show that BBB opens for low/high weight molecules and nanocarriers after exposure of loud music/sound of 90 dB and 100 dB (regardless its frequency) as being easily produced by MP3/MP4 players, kitchen appliances, loudspeakers at concerts. The role of sound, sound-induced stress and molecular mechanisms behind is discussed in the framework of BBB opening as an informative platform for a novel fundamental knowledge about the nature of BBB and for the development of a non-invasive brain drug delivery technology.

Social aspects of music/sound-induced opening of BBB provide completely new information about noise and healthy life conditions that will stimulate new research in this field.

---

The blood-brain barrier (BBB) is a highly selective barrier, which is formed by microvascular endothelial cells surrounded by pericytes and perivascular astroglia. It controls the penetration of blood-borne agents into the brain, or the release of metabolites and ions from the brain tissue to blood. Therefore, BBB plays a vital role in the central nervous system (CNS) health protecting the brain against pathogens and toxins. Although this protective mechanism is essential for normal functioning of CNS, it also creates a hindrance to the entry of drugs into the brain. Among 7,000 drugs, which are registered in the Comprehensive Medicinal Chemistry database, only 5% can successfully treat neurological diseases because 100% of large-molecule pharmaceutics (antibodies, recombinant proteins, gene therapeutics) and even smaller ones do not cross BBB^1,2^. In this context, it is not surprising that CNS diseases account for 28% of the total burden of all diseases^3^. This is the reason why approaches for reversible overcoming BBB have received significant attention in the last four decades. Currently, there are 13,443 publications registered in PubMed containing “brain drug delivery” in the title. Historically, the first 100 years of study of BBB (1880-1980) established the anatomical structure and physiology of BBB. The next 30 years were focused on the understanding of mechanisms underlying the BBB phenomenon. Now we are in the era for the development of approaches for brain drug delivery. Indeed, currently over 70 different methods were suggested for overcoming BBB including physical, chemical and biological approaches^4–6^. Nevertheless, these methods are not widely applied in daily clinical practice due to many reasons including invasiveness (e.g., photodynamic opening of BBB that requires skull opening)^7^, challenge in performing (intra-arterial injection of mannitol that only few specialists in clinics can usually do)^8^, limitation of drug concentration (intranasal drug delivery)^9^ or small area of treatment (only 1-3 mm at usage of focused ultrasound that opens BBB with additional using of micro-bubbles)^10^. All these methods require further studies to improve the reproducibility and technological robustness as well as quantitative evaluation of BBB permeability.

Despite the fact that reversible BBB breaking is a challenging and highly complicated problem for drug brain delivery, there are many acute and chronic pathological states when BBB opens itself: dementia^11^, pain^12^, hypertension^13^, rheumatoid arthritis^14^, diabetes^15^, or stroke^16^ but this increased BBB permeability in currently uncontrollable.

In general, BBB integrity depends on the appropriate functioning of tight junction machinery (paracellular permeability) and numerous transporters (transcellular permeability). Though several mechanisms controlling BBB integrity have been proposed, mechanisms underlying opening of BBB itself and influences of daily factors on BBB permeability remain unknown. The limitation of knowledge about the nature of BBB is a main factor for the slow progress in the effective therapy of CNS and the development of non-invasive methods for overcoming of BBB that can be applied in daily clinical practice.

In this study we demonstrate for the first time that such factors as loud music and specific sound, which we can meet in daily life, opens safely and reversibly BBB. Our *in vivo* and *ex vivo* results obtained in mice clearly show sound-induced opening of BBB to both low and high weight molecules as well as to nanocarriers – liposomes (100 nm-size). These results are confirmed by standard tests used for the assessment of BBB structural and functional integrity. We present in detail the music/sound impact on BBB and discuss mechanisms underlying sound-related opening of BBB and its recovery.

## Results

### I. Effects of loud music and sound on BBB permeability (ex vivo and in vivo results)

We start with the study of effects of loud music and sound on BBB permeability using classical tests with spectrofluorimetric analysis of the Evans Blue dye (EBD)-albumin complex (68.5 kDa) extravasation and confocal imaging of fluorescein isothiocyanate-dextran (FITC-dextran, 70 kDa) leakage, which are effective for the study of BBB permeability to high-weight molecules. ^17,18^

The leakage of tracers was determined in two main groups where mice have been exposed to: 1) music, Scorpions “Still Loving You”, and 2) only sound. These two groups were divided into four sub-groups: 1) control, no music/sound; 2, 3 and 4 in 1, 4 and 24h after music/sound exposure, respectively. The intensity of music/sound was 100, 90 and 70 dB; the music frequencies - 11-10,000 Hz and frequency of sound – 370 Hz.

Figure 1-I shows the music/sound-induced increase in BBB permeability to the EBD-albumin complex. One hour after music/sound exposure (100 dB, regardless frequency), the leakage of EBD was increased 17.3-fold (music), p<0.001 and 18.6-fold (sound), p<0.001 vs. the control group in all mice (100%), respectively. We found similar results in mice, that have been exposed to music/sound (90 dB): the leakage of EBD was increased 18.0-fold (music), p<0.001 and 14.6-fold (sound), p<0.001 vs. the control group in 73% of mice (11 of 15, music) and 60% of mice (9 of 15, sound). However, no changes in BBB permeability were observed when we applied a music/sound of 70 dB. The most important finding was that in 4h and 24h after loud music/sound exposure, BBB permeability to EBD was similar to the normal state (Table 1 in SI).

**Figure 1.**
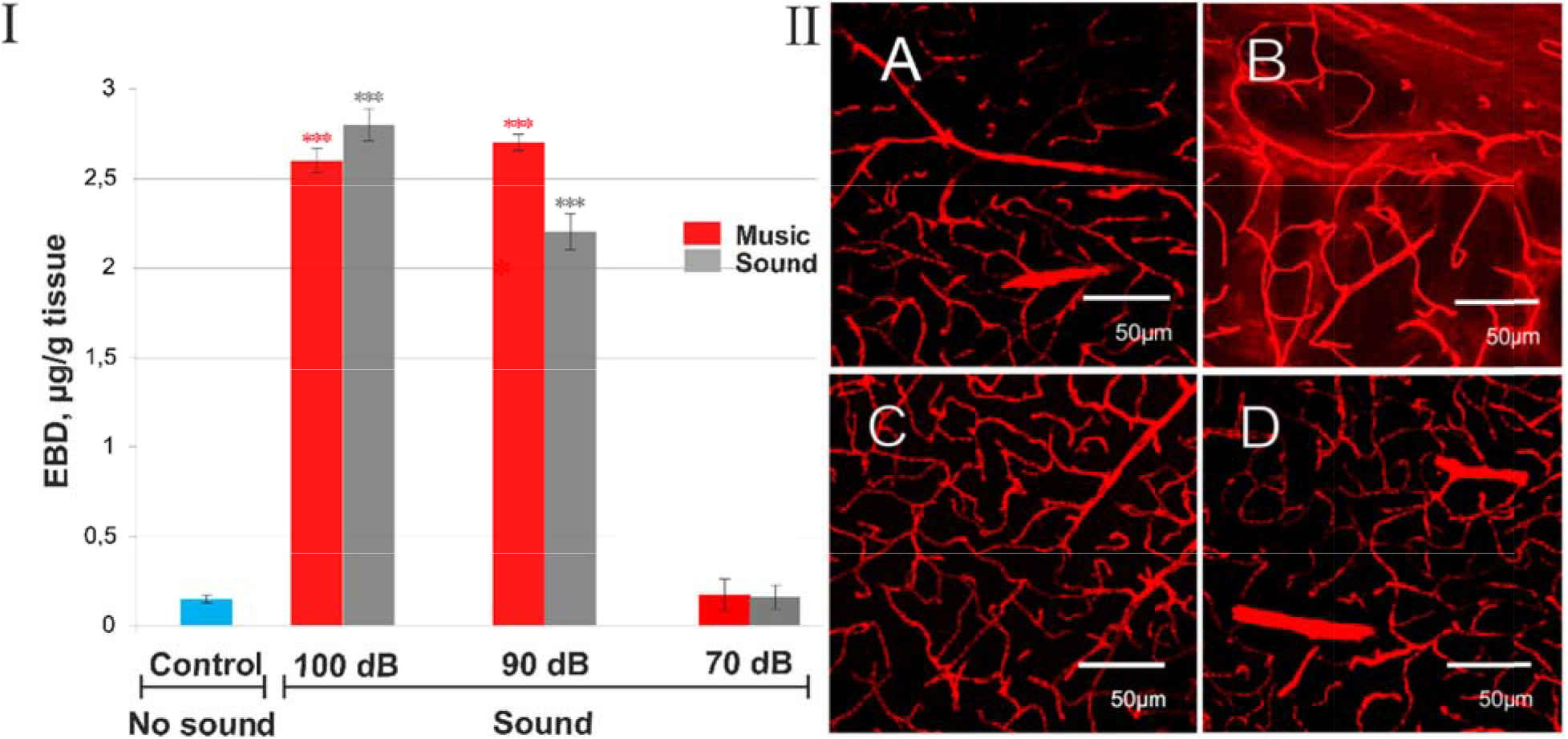
The *ex vivo* results of music/sound-induced opening of BBB. I - *The level of EBD in μg/g of mouse brain tissue before and 1h after music/sound impact.* Spectrofluorimetric assay of EBD extravasation into the brain parenchyma shows an increase in BBB permeability to EBD in 1h after the music/sound exposure. The different level of loud music/sound (100, 90 and 70 dB, regardless frequency) was accompanied by similar changes in BBB permeability to EBD in majority number of mice subjected to music/sound of 90 dB and all mice, that have been exposed to music/sound of 100 dB. No changes in BBB permeability to EBD were observed in mice subjected to sound of 70 dB. The blue color – the control group; red color – music; grey color – sound; *** - p<0.001 vs. the control group (no sound), n=15 in each group; II - Confocal imaging of BBB permeability to FITC-dextran 70 kDa in mice subjected to loud sound (100 dB, 370 Hz): A - No extravasation of FITC-dextran 70 kDa before sound exposure; B - Strong extravasation of FITC-dextran 70 kDa in 1h after sound exposure (defined as red clouds around a group of microvessels); C and D - No extravasation of FITC-dextran 70 kDa in 4h and 24h after sound exposure reflecting recovery of BBB permeability (n=10 in each group). Approximately 10 slices per animal from cortical and subcortical regions (excepting hypothalamus and choroid plexus where BBB is leaky) were imaged.

Thus, this series of experiments demonstrates that loud music/sound (100 dB) most effectively opens BBB to the EBD-albumin complex (68.5 kDa) in all tested mice and these changes are similar either for music or sound. Therefore, to keep a standardization of experimental conditions, in further steps of experiments, we used sound only with a single level and frequency (100 dB, 370 Hz).

*Ex vivo* confocal microscopy revealed the accumulation of FITC-dextran 70 kDa outside of brain capillaries in 1h after sound exposure. Extravasation of FITC-dextran was determined as a clear fluorescence visualized outside the vessel walls (Figure 1, II-B). No extravasation of FITC-dextran was observed in the control group (no sound), in 4h and 24h after the sound exposure (Figure 1, II-A, C, D,).

Collectively, our *ex vivo* results uncover that loud music/sound 90 and 100 dB (regardless frequency) reversibly opens BBB in mice with a similar degree of BBB permeability to high weigh molecular substances such as the EBD-albumin complex 68.5 kDa and FITC-dextran 70 kDa.

*In vivo* two-photon laser scanning microscopy (2PLSM) of the FITC-dextran leakage confirmed our results obtained by *ex vivo* confocal imaging. Figure 2, I A-D demonstrates the FITC-dextran leakage in 1h after sound exposure. Figures 2, I E-G illustrates the FITC-dextran extravasation kinetics that reached a plateau at ~7.5 min after its injection in 1h after sound exposure and returned to the normal state in 4h after sound impact. We observed again small the FITC-dextran extravasation with a plateau to ~12.5 min in 24h after sound exposure. However, these changes were statistically not significant and could be explained by inflammation processes developing after craniectomy. Note, in an intact brain the FITC-dextran extravasation was not observed.

**Figure 2.**
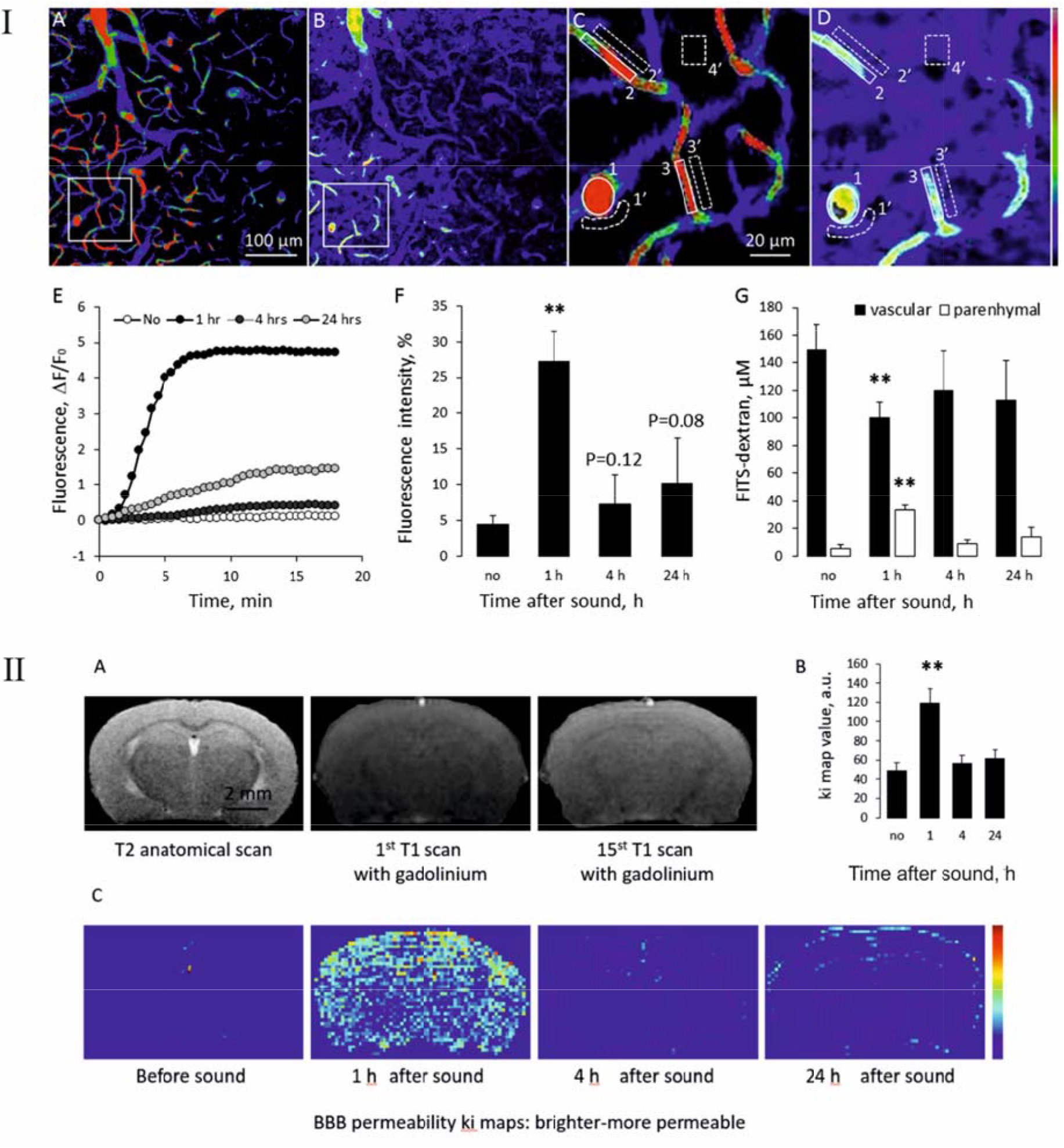
The *in vivo* results of sound-induced BBB opening. I - *In vivo* two-photon laser scanning microscopy of BBB permeability for FITC-dextran 70 kDa in mice subjected to sound (100 dB, 370 Hz): A - The representative micrographs of mouse parietal cortex microvasculature in 1h after sound exposure just after injection of FITC-dextran 70 kDa; B - The same region 20 min after FITC-dextran injection; C - Cortical region at higher magnification from (A) depicting quantification of microvascular permeability. Blood vessel area (1-3) and interstitial space (1’-4’) were manually defined and fluorescence intensity in each area was evaluated using the ImageJ software; D - The same area at higher magnification from (B); E - Time course of FITC-dextran extravasation showing an increased BBB permeability in 1h after sound exposure. Data are presented as mean ± SEM, n = 7; F - The percentage fluorescence intensity in perivascular area (calculated as % of the fluorescence in the vascular area) reflecting maximum FITC-dextran leakage in 1h after sound. Data are presented as mean ± SEM, n = 7, **p < 0.01; G - The estimation of FITC-dextran concentration in the brain parenchyma and microvessels in 20 min after injection. Data are presented as mean ± SEM, n = 7, **p < 0.01. II -MRI analysis of BBB permeability in mice subjected to sound: A - The MRI data representing T2 (anatomical scan) and 1^st^ and 15^th^ rapid T1 MRI scan after Gd-DTPA injection; B - The *Ki* value:s show (arbitrary units) rate of changes in the MRI signal intensity, reflecting essential Gd-DTPA leakage in 1h after sound with rapid recovery of BBB. Data are presented as mean ± SEM, n = 7, **p < 0.01; C - The *Ki*-maps from the same mice at different time points showing maximal BBB permeability to Gd-DTPA in 1h after sound exposure (brighter is more permeable).

In the next step, we analyzed effects of sound on BBB permeability to low weight substances. We performed a magnetic resonance imaging (MRI) analysis of gadolinium-diethylene-triamine-pentaacetic acid (Gd-DTPA, 938 Da) leakage by using a custom-made sequence (Ki values) for a post hoc evaluation of changes in MRI signal intensity.^19^ The anatomical scan of the mouse brain with T2 and rapid T1 for Gd-DTPA analysis is presented in Figure 2, II-A. The data given in Figure 2, II-B indicate a statistically significant (p<0.01) increase in BBB permeability to Gd-DTPA in all regions of the brain in 1h after sound exposure. Note, the Ki map values reached 185% (119.3) arbitrary units (p<0.01) only in 1h after sound exposure, while in 4h and 24h, essential changes in the Ki values were not observed, compared with the control group (no sound). Similar results obtained from Ki maps of BBB permeability and presented in Figure 2, C, reflect the maximal Gd-DTPA leakage only in 1h after sound exposure (brighter – more permeable).

Thus, our *in vivo* results confirmed *ex vivo* data that BBB is opened for high and low weight molecules in 1h after sound exposure (100 dB, 370 Hz) followed by rapid recovery of BBB permeability.

### II. BBB permeability to nanomaterials (GM_1_-liposomes)

To evaluate the applied significance of the sound-induced opening of BBB, we next performed an *ex vivo* and *in vivo* study of BBB permeability to fluorescently labelled 100-nm liposomes supplemented with ganglioside GM_1_ (GM_1_-liposomes; Figure 3, A, B in SI) as good candidates for brain drug delivery systems and therapy of brain diseases. Previously, ganglioside GM_1_ having the ability to stabilize liposomes in the circulation was also shown to contribute to their transport across BBB^20^.

Figure 3 (I-II) shows the results of the confocal (ex vivo) analysis of BBB permeability to GM_1_-liposomes. One hour after sound, GM_1_-liposomes were observed outside of endothelial cells and between astrocytes labeled by SMI-71 and GFAP, respectively. There was no extravasation of GM_1_-liposomes before as well as in 4h and 24h after sound exposure.

**Figure 3.**
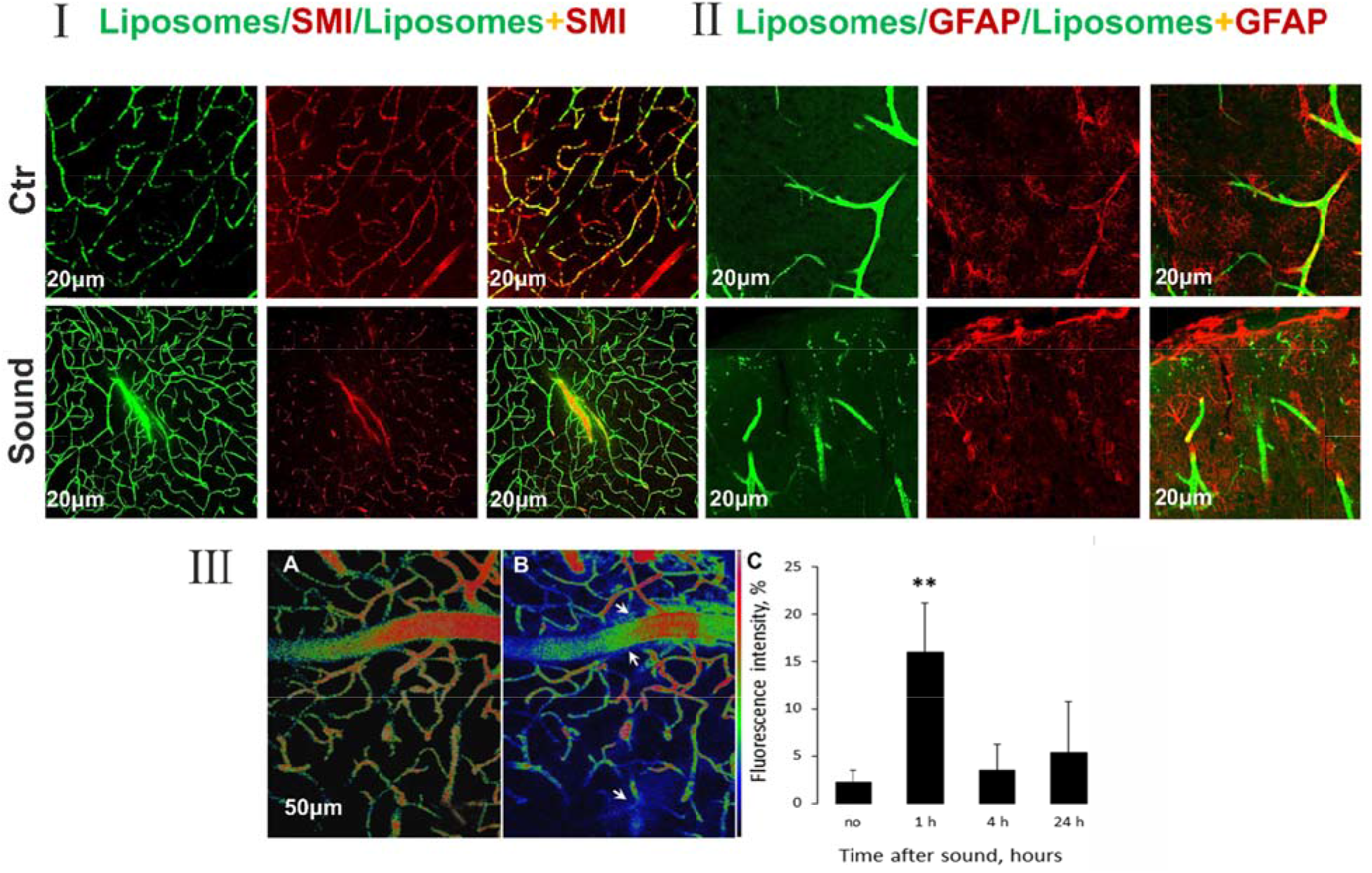
The sound-induced increase in BBB permeability to GMļ-liposomes. I-II - Confocal microscopy of BBB permeability for GM_1_-liposomes 100 nm (green fluorescence) in mice (n=10) in 1h after sound exposure (100 dB, 370 Hz) using specific markers of BBB: I - red color is a marker of cerebral endothelial cells (SMI-71); II – red color is a marker of astrocytes (GFAP). There is extravasation of GM_1_-liposomes 100 nm (defined as a green cloud around cerebral vessels) in 1h after the sound exposure. No extravasation of GM_1_-liposomes 100 nm under the normal state (Ctr) is seen. Approximately 10 slices per animal from cortical and subcortical regions (excepting hypothalamus and choroid plexus where BBB is leaky) were imaged. III - *In vivo* two-photon laser scanning microscopy of BBB permeability for GM_1_-liposomes (100 nm) in mice subjected to sound (100 dB, 370 Hz): A - Representative micrographs of mouse parietal cortex microvasculature in 1h after sound exposure just after intravenous injection of GM_1_-liposomes; B - The same region 20 min after injection. The arrows show the leakage area in perivascular tissue; C - The fluorescence intensity (calculated as % of the fluorescence in the vascular area) in the perivascular area reflecting the maximum BBB leakage in 1h after sound exposure. Data are presented as mean ± SEM, n = 7, **p < 0.01.

The *in vivo* 2PLSM of fluorescent GM_1_-liposomes leakage confirmed our *ex vivo* confocal data. Figure 3, III demonstrates the increase of perivascular parenchyma fluorescence to 15.9 ± 5.2% in 1h (p<0.01) after sound exposure suggesting opening of BBB for GM_1_-liposomes. There were no statistically significant changes in BBB permeability to GM_1_-liposomes in 4h and 24h after sound action. In an intact brain, extravasation of GM_1_-liposomes was also not observed.

Thus, our *ex vivo* confocal data with using of BBB markers and *in vivo* 2PLSM imaging suggest that GM_1_-liposomes 100 nm-size effectively cross BBB in 1 h after sound exposure.

### III. Mechanisms underlying sound-induced opening of BBB

#### i. Sound opens BBB directly and via stress

To answer the question, whether sound opens BBB directly or via stress, we further analyzed BBB permeability to the EBD-albumin complex in anesthetized mice to prevent influences of stress stimuli. Additionally, we studied sound effects on the BBB integrity in *in vitro* experiments using measurement of trans-endothelial electrical resistance (TEER)^21^.

The BBB permeability to the EBD-albumin complex was 6.7-fold lesser in anesthetized mice compared with mice affected by sound without anesthesia (0.38±0.03 vs. 2.55±0.07, p<0.001). This fact emphasizes that stress is an important factor for sound-induced opening of BBB. However, even under anesthesia sound increased EBD leakage up to 2.5-fold (0.38±0.03 vs. 0.15±0.01, p<0.001).

Before measuring of TEER, we established an optimal time of sound effects on BBB in *in vitro* model by studying the appearance of blebbing brain cells exposed to sound during 5, 10, 20, or 40 min (Figure 4, SI). We found that 5 and 10 min of sound exposure was associated with the minimal induction of cell membrane blebbing in brain cells. Therefore, this regimen was further used for the assessment of TEER. Figure 4B shows a significant decrease in TEER 5-10 min after sound exposure reflecting opening of BBB and confirming direct sound effects on BBB.

**Figure 4.**
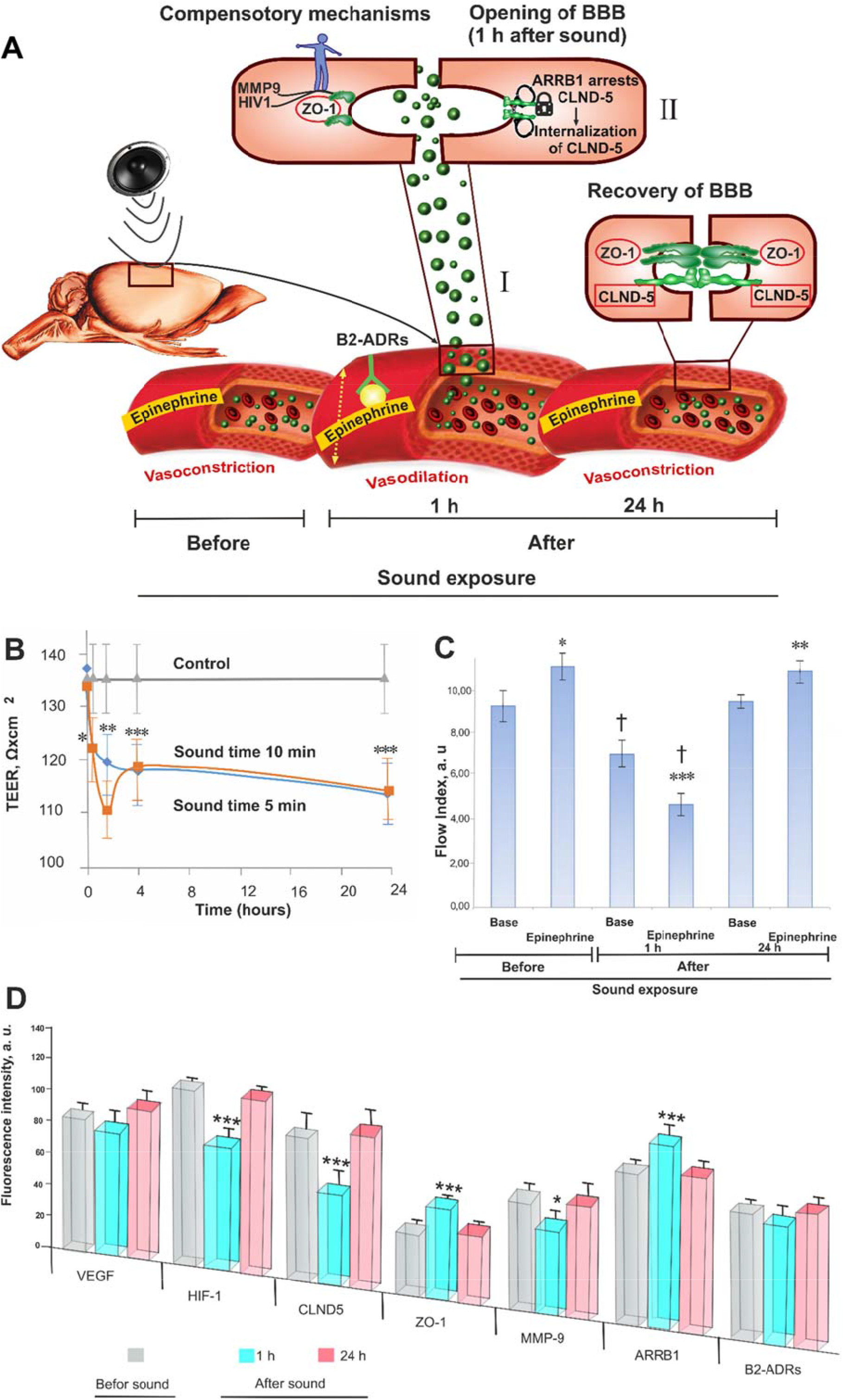
Mechanisms underlying sound-induced opening of BBB. A – The scheme of hypothesis explaining mechanisms underlying sound-induced opening of BBB: I – The opening of BBB is associated with a decrease in the CBF reflecting vasodilation that might be stimulated by epinephrine-related activation of B2-ADRs; II - The ARRB1 is a co-factor in signaling pathways of activation of B2-ADRs. The increase in the expression of ARRB1 is accompanied by a decrease in the expression of CLND-5 that might be a reason of its “arrest” (loss of surface in TJ space) by ARRB1 that contributes opening of BBB; III - The compensatory mechanisms such as decrease in the expression of HIV-1 and MMP-9 lead to the rapid recovery of BBB permeability. B – The sound-induced changes in TEER in BBB model *in vitro*. One hour after sound exposure (during 5 and 10 min, n=6 in each group) TEER is significantly decreased (p<0.05) suggesting opening of BBB; C - LSCI monitoring of epinephrine effects (iv.) on the CBF at 7 mice: epinephrine causes an increase in the CBF when BBB is closed (in the normal state and 24h after sound exposure) and a decrease in the CBF when BBB is opened (1h after sound exposure). * - p<0.05; ** - p<0.01; *** - p<0.001 vs. the basal level of CBF (before epinephrine injection); † - p<0.05 vs. the control group (before sound). D – The expression of molecular factors of regulation of endothelial permeanility in the control group (before sound exposure, 1h and 24h after sound (n=7 for each group): opening of BBB (1h after sound exposure) is associated with a decrease in the expression of CLND-5 (p<0.001), MMP-9 (p<0.05), HIV-1 (p<0.001), and an increase in the expression of ZO-1 (p<0.001), ARRB1 (p<0.001) but no changes in B2-ADRs and VEGF. The expression of the analyzed molecular factors is returned to the normal state in the next day after sound exposure. * - p<0.05; ** - p<0.01; *** - p<0.001 vs. the control group (before sound).

#### ii. Stress-induced humoral and hemodynamic responses after sound-induced opening of BBB

In the previous set of experiments, we uncovered that stress is an important factor in sound-induced opening of BBB. To study the role of sound stress in a reversible opening of BBB, here we analyzed the plasma level of epinephrine as the key stress hormone using enzyme immunoassay and the cerebrovascular effects of epinephrine using laser speckle imaging.

It was expected that sound stress caused an increase in the plasma level of epinephrine up to 7.0-fold vs. the normal state (23.1±2.7 ng/ml vs. 3.3±0.9 ng/ml, p<0.001). One hour after sound, the level of epinephrine decreased and reached 8.7±1.9 ng/ml (p<0.001).

Our results showed that during BBB opening (1h after sound exposure) the basal CBF was decreased by 24±3% (7.0±0.6 a.u. vs. 9.2±0.7 a.u., p<0.05) compared with the control group (Figure 4C). On the next day when BBB was closed, the basal CBF completely recovered and was similar to the normal state (9.4±0.3 a.u. vs. 9.2±0.7 a.u.).

We found that in the control group (before sound exposure) epinephrine caused an increase in the CBF by 19±2% (11.0±0.6 a.u. vs. 9.2±0.7 a.u., p<0.05) suggesting the vasoconstriction and increase of vascular tone. During BBB opening (1h after sound exposure), epinephrine induced an opposite vascular response – a decrease in the CBF by 33±5% (4.7±0.5 a.u. vs. 7.0±0.6 a.u., p<0.001) reflecting vasorelaxation and a decrease of vascular tone. It is important to note that in 24h after sound exposure the CBF responses to epinephrine recovered and were typical to the normal state, i.e. epinephrine caused an increase in the CBF by 14±3% (10.8±0.5 a.u. vs. 9.4±0.3 a.u., p<0.01).

#### iii. Molecular factors involved in sound-induced opening of BBB

Above, we showed that opening of BBB was accompanied by a decrease in CBF reflecting vasorelaxation that was enhanced by the injection of epinephrine.

The epinephrine causes the endothelial relaxation via activation of vascular beta2-adrenoreceptors (B2-ADRs)^22^. The beta-arrestin-1 (ARRB1) is a membrane co-factor of activation of B2-ADRs playing an important role in vascular responses to stress.^23,24^ Taking into account these facts, we hypothesized the possible role of the adrenergic mechanism of vasorelaxation in sound-induced opening of BBB.

Therefore, we next studied the expression of vascular B2-ADRs in the brain and ARRB1 in the brain and blood.

Furthermore, we evaluated the expression of TJ (CLND-5 and ZO-1) as well as molecular factors (VEGF, HIV-1, MMP9) playing an important role in the regulation of the endothelial permeability.^25–27^

Figure 4D shows that a sound-induced opening of BBB was associated with a decrease in the expression of CLND-5 (p<0.001), MMP-9 (p<0.05), HIV-1 (p<0.001), and an increase in the expression of ZO-1 (p<0.001), ARRB1 (p<0.001) but no changes in B2-ADRs and VEGF. It is important to note that the expression of the analyzed molecular factors was recovered to the next day after sound exposure when BBB was closed.

#### iv. Sound opens BBB safely

To answer the question, how sound affects the brain and its functions, we studied morphological changes in the brain tissues using histological methods and assessed hearing and cognitive impairment using the *Fear conditioning* (FC) test^28–30^.

The histological examination did not reveal any changes in the brain tissues and vessels after the temporal sound-related opening of BBB (Figure 7, SI).

Applying the method of TUNEL *in vitro* experiments, we found that sound did not induce excessive apoptosis in the rat brain cortex in 2.5 and 5 min of sound exposure (Figure 6, SI).

We found that loud sound did not affect hearing or cognitive functions (Figure 5, SI). Mice demonstrated the same total freezing time either in the control (before sound) or experimental groups (1h and 24h after sound exposure 100 dB): 190±27 s, 213±24 s, 176±3 s, respectively (one-way ANOVA, p=0.4181). The analysis of the time segments revealed no significant difference in the freezing time between the control and experimental groups: 28±1 s, 28±1 s, 27±2 s, respectively (one-way ANOVA, p=0.6833), or in the freezing latency: 0.9±0.6 s; 3.4±2.3 s, 1.6±0.6 s, respectively (one-way ANOVA, p=0.8724).

**Figure 5.**
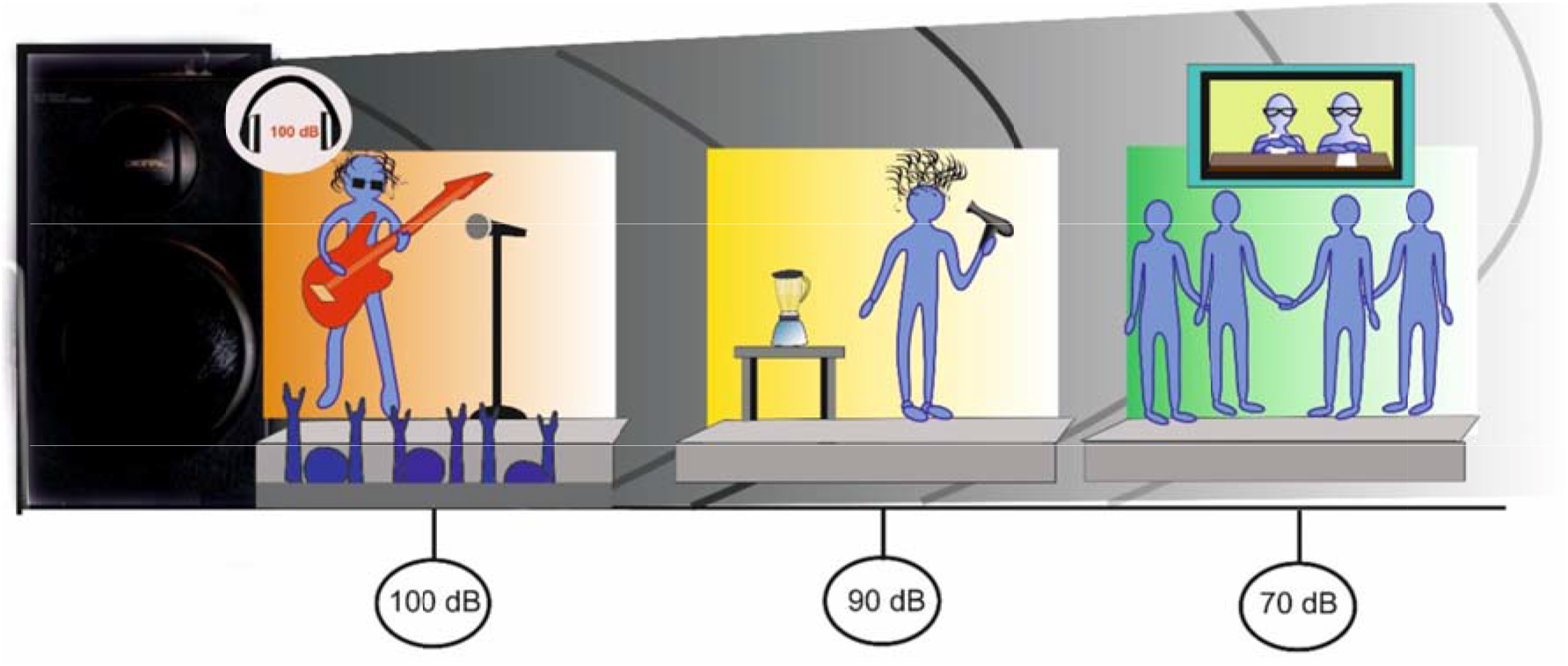
Examples of loud music and sound in our life. Sound pressure of 100 dB: Some MP3/MP4 players produce sound of 100-110 dB during listening music with earphones; we have high energy sound (100-110 dB) during listening to music at a rock concert; monkeys or dogs generate very high noise levels (100-110 dB) by rattling and banging on their laboratory cages;^30–32^ 90 dB is typical for daily life: hair dryer, kitchen blender, or food processor and 70 dB: TV, radio, group conversation.^31, 32^

Thus, loud sound (100 dB, 370 Hz) did not affect hearing, cognitive functions or behavior in mice.

## Discussion

Our *in vivo* and *ex vivo* results, for the first time, clearly show that such natural factors as loud music/sound (90-100 dB, regardless frequency), which humans and animals meet in daily life^31–33^, reversibly opens BBB for low and high weight molecules as well as to 100-nm liposomes for a short period followed by a quick recovery. The novel aspects of mechanisms underlying the phenomenon of sound-induced transient opening of BBB were also described.

The use of different frequencies and intensities of sound to deliver drugs into the brain has been shown over 40 years ago. There are numerous studies using infrasound or ultrasound for transient BBB opening.^10, 34–38^ At present, infrasound (8 Hz, 90-120 dB, duration – 2h) effectively opens BBB but induces significant brain injuries due to increased intracellular calcium concentration, induction of apoptosis and high expression of heat shock protein 70.^34,35^ The currently focused ultrasound is one of the most promising methods to enhance drug penetration into the brain by disruption the TJ between capillary endothelial cells.^10, 36–38^ However, this method remains largely experimental and continues to make progress toward becoming a clinical tool. Also, a range of substantial limitations of focused ultrasound such as tiny window (1-3 mm),^10^ low penetration (100-200 μm),^39^ no effectiveness without gas microbubbles,^39^ appearance of apoptosis and ischemic areas,^40^ hemorrhages,^41^ and activation of sterile inflammation^42^ prevents its future progression toward a clinical application.

It should be emphasized that no successful clinical trial has been reported yet. The main reason for the slow progress in drug brain delivery is a lack of knowledge about the nature of BBB.

It is well known that BBB opens itself in different acute and chronic pathological states^11–16^. In our study we demonstrated, for the first time, that BBB is opened also in not a pathological condition. Taking into account the fact that different types of sound induce BBB disruption^10, 34–38^, here we focused on the study of effects of natural factors such as loud music/sound on BBB permeability.

In this research, we addressed the following questions.

i. How does sound open BBB? We found that sound opens BBB after 2h of exposure (60 s – sound; 60 s pause) with a latent period - in 1h after sound action followed by complete recovery of BBB permeability in 4h after sound influence. We also demonstrated similar results by using of the photodynamic opening of BBB^7^. It might be proposed that sound might open BBB by “All-or-none law” firstly established by the American physiologist Henry Pickering Bowditch in 1871. According to him the strength by which nerves or muscles respond to a stimulus is independent of the strength of the stimulus if it exceeds the threshold potential. So, the sound of 90 dB effectively opened BBB. A further increase in the sound level to 100 dB was not accompanied by a more pronounced degree of BBB permeability – “All” and the sound 70 dB did not open BBB – “Nothing”. In *ex vivo* experiments using spectrofluorimetric assay and confocal imaging, we showed sound-induced opening of BBB for high weight molecules such as the EBD-albumin complex 68.5 kDa, FITC-dextran 70 kDa as well for 100 nm-GM_1_-liposomes that was confirmed in *in vivo* experiments using 2PLSM. Using MRI, we demonstrated BBB opening for low weight molecules such as Gd-DTPA (928 Da). These results suggest the high efficiency of sound to reversibly open BBB to low/high weight molecules and nano-carriers such as 100nm-GM_1_-liposomes.
ii. Does sound open BBB safely? We demonstrated the efficiency of loud sound (90 and 100 dB) to open BBB in a safe regime (60 s – sound; 60 s – pause during 2h) according to the rules for safe listening.^31^ Assessment of apoptosis, brain histology and neurobehavioral pattern did not reveal excessive apoptosis, morphological alterations in the brain tissues and vessels, or hearing loss and cognitive impairment in mice in 1h after sound exposure when BBB was opened and in 4h/24h after sound exposure when BBB integrity was fully restored. The reversibility of sound-induced BBB breakdown might be linked to the activation of the brain drainage and clearing systems via the meningeal lymphatic pathway after BBB opening ^43,44^. In our recent work, we discovered an important role of the meningeal lymphatic systems in the brain clearing from substances, which permeate from blood into the brain parenchyma through opened BBB induced by sound or photodynamic effects.^43, 44^
iii. Does sound open BBB directly or via stress? We analyzed BBB permeability to the EBD-albumin complex under anesthesia, which blocks stress influences and in *in vitro* experiments using a model of BBB. The results showed that anesthesia significantly reduced the BBB breakdown mediated by an acoustic action. We used gas anesthesia (isoflurane) that minimally affects BBB.^45^ These results indicate the important role of stress in opening of BBB induced by a loud sound. However, in *in vitro* experiments using the model of BBB, we observed that sound caused a decrease in TEER, thereby suggesting direct BBB disruption^20^. Also, we observed that even under anesthesia BBB remained open to the EBD-albumin complex. These data clearly show that both factors – sound itself and stress play an important role in opening of BBB.
iv. What mechanisms are responsible for sound-induced opening of BBB? We analyzed systemic and molecular mechanisms underlying the phenomenon of sound-induced opening of BBB. At the systemic level, sound stress was associated with a high plasma concentration of epinephrine that decreased in the post-stress period when BBB was opened (1h after sound exposure). The elevated level of stress hormones, including catecholamine, induced by sound, has also been shown in humans and animals as discussed in Turner et al.^32^ At the hemodynamic level, sound-induced opening of BBB caused the conversion of vascular effects of epinephrine (reduction of CBF), while BBB was closed (in the control group or 24 h after sound exposure) epinephrine induced an increase in CBF. The epinephrine-induced decrease in CBF after sound stress might be a reason of vasodilation via stimulation of B2-ADRs that are responsible for a smooth muscle relaxation^22^. In the post-stress period, the activation of B2-ADRs is predominant due to their high sensitivity to the low level of epinephrine^46^ that we observed after sound exposure. However, this hypothesis is applicable at the artery/arteriole level of brain microcirculation, but at the capillary level where smooth muscle cells are absent, a possible activation of adrenergic receptors expressed on astroglial cells or pericytes^47,48^ should be taken into consideration. We did not reveal any changes in the expression of vascular B2-ADRs in mice underwent sound exposure, but we found an increase in the expression of ARRB1, which is a signaling adaptor protein and marshal in trans-membrane scenarios of activated B2-ADRs under normal as well as pathological conditions^23,24^.

Our results revealed that the elevated expression of ARRB1 was associated with a decrease in the expression of CLND-5 as the main component of BBB integrity.^49^ We suppose that the ARRB1 “arrests” the expression of CLND-5 by stimulating its internalization. The loss of surface of CLND-5 in space between endothelial cells can be one of the mechanisms underlying sound-induced opening of BBB. Our hypothesis is very close to the mechanism of BBB disruption via ARRB2-mediated internalization of VE-cadherin induced by VEGF.^50^ The elevated expression of ZO-1 during sound-induced opening of BBB on the background of lowered expression of CLDN-5 might also suggest about disordered interactions of TJ proteins.

The sound-induced opening of BBB is reversible. We demonstrated that BBB was closed in 4h after sound exposure. We suppose that it might be related to compensatory mechanisms via reduced expression of MMP-9 and HIV-1 that control endothelial permeability and survival.^26,27^ It is well known that a high expression of these factors is accompanied by the BBB breakdown. It is interesting to note that if loud sound is used in a non-safe regime (sound trauma, 120 dB during 3h), it is accompanied by a significant increase in the expression of HIV-1 and VEGF with elevated endothelial permeability and pericytes alterations.^51^

In sum, the BBB breakdown induced by sound exposure causes a reversible disorganization of the TJ machinery in affected cerebral endothelial cells (Figure 4A and movie in SI).

## Conclusion

We show that loud music and sound effectively open BBB to high and low weight molecules as well to nano-carriers (100 nm-GM_1_-liposomes). The loud sound directly or via stress-mediated mechanisms acts at the BBB tight junction machinery.

Loud music and sound are factors, which we have in our daily life during listening of MP3/MP4 players using earphones or kitchen techniques or attending music concerts etc. These natural factors open BBB safely without morphological changes in the brain tissues or vessels, presentence of brain apoptotic cells, hearing loss and cognitive impairment. Despite the fact that music/sound opens BBB in many regions of the brain, i.e. in a non-specific manner, we strongly believe that this method has a high potential for clinical applications as an easily used, non-invasive, low cost, labeling free, perspective and completely new approach for the treatment of brain diseases (i.e. neurodegenerative disorders).

The fact that loud music/sound opens BBB even in an ordinary (safe) regime is socially important and should be considered in daily life.

## Methods

### Subjects

The experiments were done on four groups of male mongrel mice (20-25 mg): (1) no sound – the control group; 2, 3 and 4 – in 1, 4 and 24 h after sound exposure, respectively. For some experiments, Wistar male rats of corresponding age been used. All procedures were performed in accordance with the “Guide for the Care and Use of Laboratory Animals.”^52^ The experimental protocols were approved by the Local Bioethics Commission of the Saratov State University (Protocol 7); Experimental Animal Management Ordinance of Hubei Province, P. R. China (No. 1000639903375); Local Bioethic Commission of the Krasnoyarsk State Medical University named after Prof. V.F. Voino-Yasenetsky (Russia); the Institutional Animal Care and Use Committee of the University of New Mexico, USA (#200247); the Institutional Animal Care and Use Committee of Shemyakin-Ovchinnikov Institute of Bioorganic Chemistry, Russian Academy of Sciences, Russia.

### Experimental design of music/sound effects on BBB permeability.

To produce the loud music (100 dB, 11-10,000 Hz, Scorpions “Still loving you” was used as an example, see Figure 1 in SI, the spectrum was calculated using free software for sound proceeding – Audacity ®) and sound (100 dB, 370 Hz) we used loudspeaker (range of sound intensity – 0-130 dB, frequencies - 63-15000 Hz; 100 V, size: 450×640×330 mm, Yerasov Music Corporation, Saint Petersburg, Russia) and sound transducer (100 dB, 7A, 12 V, Auto VAZ PJSC, Tolyatti, Russia) using consequence of: 60 s – sound on, then 60 s – sound off during 2h. The sound level was measured directly in a cage of animals using sound level meter (Megeon 92130, Russia, see Figure 2 in SI).

According official information of American Speech-Language-Hearing Association^31^ we selected three types of sound pressure: 1) extremely loud - 100 dB, which is typical for some MP3/MP4 players producing sound of 100-110 dB during listening music with earphones; 2) very loud - 90 dB, which we can listen using hair dryer, kitchen blender, or food processor; 3) moderate - 70 dB – radio or TV, group conversation^31, 32^ (Figure 5).

The mice have hearing with frequency sensitivity up to the ultrasonic range (> 80,000 Hz)^33^, far beyond the frequencies of music/sound (11-10,000 Hz), which we used in experiments. Nevertheless, to make experimental condition closer to real life, we selected two types of sound – loud music 70-100 dB with frequencies in the range of 10-10,000 Hz and maximal intensity around 100 Hz, and loud sound 70-100 dB with a single frequency 370 Hz (Figure 1, SI).

Duration of sound was 2h with intermittent mode: 60 s – sound and 60 s - pause. Our choice was related to the fact that sound of 85 dB can lead to hearing loss if we listen for it continuously for more than 8 h^31^. The safe listening time decreases in a half for every 3 dB rise of sound levels over 85 dB. Thus, for the sound of 91 dB, safe listening time is down to 2h and for sound 100 dB – 15 min. However, it was no effect found on the BBB when using the sound of 100 dB during 15 min (data not presented). Therefore, we selected 2h exposure for all sound intensities (100-90-70 dB) but to provide safety an intermittent mode of sound application was used.

### Spectrofluorimetric assay of EBD extravasation

The mice were placed on a heating platform to maintain body temperature during all the steps of the experiments. Polyethylene catheters (PE-10 tip, Scientific Commodities INC., Lake Havasu City, Arizona) were inserted into the right femoral vein for EBD (Sigma Chemical Co., St. Louis, MI, USA) intravenous injection in a single bolus dose (2 mg/25 g mouse, 1% solution in physiological 0.9% saline). The EBD circulated in the blood for 30 min in accordance with the recommended protocol.^53^ The implantation of the catheter was performed under the inhalation anesthesia (2% isoflurane, 70% N_2_O and 30% O_2_). At the end of the circulation time, mice were killed by decapitation, their brains, liver and blood were quickly collected and placed on ice (no anti-coagulation was used during blood collection).

The detailed protocol of EBD extraction and visualization was published by Wang et al.^53^ Briefly, the blood samples were centrifugated at 10.000 x g (for 20 min). Prior brain removal, the brain was perfused with 0.9% saline to wash out remaining dye in the cerebral vessels. The isolated brain were cut into small pieces and incubated with 0.9% saline (1:3) for 60 min to enable the soluble substances to dissolve. Then the solutions were centrifugated at 10.000 x g for 10 min to sediment the non-dissolved parts. The supernatants that contained the plasma or the brain solutes were treated by 50% trichloroacetic acid - TCA (1:3), centrifugated again at 10.000 x g for 20 min to remove precipitated molecules. The final supernatants were incubated in 95% ethanol to increase the optical signals for the spectrofluorometric assay (620 nm/680 nm, Agilent Cary Eclipse, Agilent, USA). The standard calibration curve was created and used for calculation of the EBD concentration (μg per g of tissues or blood).

The leakage of EBD was determined in two main groups where mice have been exposed to: 1) music, Scorpions “Still Loving You”, and 2) only sound. These two groups were divided into four sub-groups: 1) control, no music/sound; 2, 3 and 4 in 1, 4 and 24h after music/sound exposure, respectively, n=15 in each group.

The effects of anesthesia (2% isoflurane at 1L/min N_2_O/O_2_ – 70:30) on the leakage of EBD was studied in two groups: (1) control, no sound (n=17); in 1h after sound exposure (n=20).

### Confocal microscopy of FITC-dextran 70 kDa extravasation

Fluorescein isothiocyanate (FITC)-dextran 70 kDa was used as an additional valuable tool to characterize the BBB permeability to high weight molecules. The BBB permeability to dextran was evaluated *in vitro* by confocal microscopy and *in vivo* by two-photon laser scanning microscopy.

The protocol for high weight fluorescein isothiocyanate (FITC)-dextran to assess BBB disruption using confocal microscopy is described in details.^18^ Briefly, given a description of the main steps of this protocol. FITC-dextran 70 kDa was injected intravenously (1 mg/25 g mouse, 0.5% solution in 0.9% physiological saline, Sigma-Aldrich) and circulated for 2 min. Afterward mice were decapitated and the brains were quickly removed and fixed in 4% paraformaldehyde (PFA) for 24 h, cut into 50-μm thick slices on a vibratome (Leica VT 1000S Microsystem, Germany) and analyzed using a confocal microscope (Olympus FV10i-W, Olympus, Japan). Approximately 8–12 slices per animal from cortical and subcortical (excepting hypothalamus and choroid plexus where the BBB is leaky) regions were imaged. The confocal microscopy of the BBB permeability performed in the groups: 1) control, no sound (n=10); 2) at 1h (n=10); at 4h (n=10); at 24h (n=10) after sound exposure.

### In vivo real-time two-photon laser scanning microscopy (2PLSM)

BBB permeability was continuously monitored by measuring the perivascular tissue fluorescence in 7 mice at different time points: before and in 1, 4 and 24h after sound exposure as described previously with some modifications.^54^ Three days before imaging, chronic optical windows (ø 3 mm) were prepared in coordinates 1-4 mm caudal and 1-4 mm lateral to bregma as we previously described.^54^

During the imaging mice were kept under inhalation anesthesia with 2% isoflurane at 1L/min N_2_O/O_2_ – 70:30. The body temperature was maintained at 37.5°C by a homoeothermic blanket system with a rectal probe (Harvard Apparatus, Holliston, MA). The brain temperature was kept at 37°C using an objective heating system with a temperature probe (Bioptechs Inc., Butler, PA). The fluorescein-labeled dextran (70 kDa, Sigma-Aldrich) in physiological saline (5% wt/vol) was injected through the tail vein (~100 μl) at an estimated initial concentration in blood serum of 150 μM.^54^

Imaging was done using Prairie View Ultima system with Olympus BX51WI upright microscope and water-immersion LUMPlan FL/IR 20×/0.50 W objective. The excitation was provided by a Prairie View Ultima multiphoton laser scan unit powered by a Millennia Prime 10 W diode laser source pumping a Tsunami Ti: sapphire laser (Spectra-Physics) tuned up to 810 nm central wavelength. Fluorescence was band-pass filtered at 510–550 nm. Real time images (512 × 512 pixels, 0.15 μm/pixel in the *x*- and *y*-axes) were acquired using time series mode of the Prairie View software (30 s interval between images, 30 min session). Between the sessions (before and in 1-4-24h after sound exposure) mice were awake and were kept at the animal facility.

In off-line analyses using the ImageJ, microvascular BBB permeability was evaluated by measuring changes in perivascular tissue fluorescence in planar images of the cortex taken 50 and 150 μm depth in 20 min after fluorescein dextran injection, as described previously.^54^ For each image, an intensity of fluorescence of ten randomly chosen regions of interest over the vessel areas and 10 corresponding perivascular brain parenchyma areas were evaluated. The obtained estimates in the interstitial space were normalized to the maximal fluorescence intensity in the blood vessels and expressed as a percentage of fluorescence intensity (modified technique from Egawa et al.).^55^ The estimated quantity of fluorescein, that leaked out of the vessel and left in the vessel for 20 min after the injection, was calculated from the initial vascular fluorescence equal to ~150 μM of fluorescein. The time courses of fluorescein leakage for each group were also plotted.

### Magnetic Resonance Imaging (MRI) for the analysis of BBB permeability

MRI was conducted on the same mice, which we used for 2PLSM, at different time intervals 0 – before and at 1, 4, 24h after sound exposure on a 7-T dedicated research MRI scanner (Bruker Biospin; Billerica, MA, USA). Signal transmission and detection were done with a small-bore linear RF coil (inner diameter of 72 mm) and a single tuned surface coil (RAPID Biomedical, Rimpar, Germany). The mice kept under inhalation anesthesia (2% isoflurane at 1L/min N_2_O/O_2_ – 70:30). Respiration and heart rate were monitored during MRI measurements, and the body temperature was maintained at 37.0 ± 0.5°C.

Anatomical T2-weighted images were acquired before each BBB permeability determination with a fast spin-echo sequence (rapid acquisition with relaxation enhancement (RARE)) (Repetition Time (TR)/Echo Time (TE) = 5,000 ms/56 ms, Field of View (FOV) = 4 cm × 4 cm, slice thickness =1 mm, slice gap (inter-slice distance) =1.1 mm, number of slices =12, matrix = 256 × 256, number of averaging = 3) as previously described.^54^

To non-invasively evaluate the BBB permeability, we used a modified dynamic contrast-enhanced (DCE)-MRI and graphical analysis of the resultant image data.^56^ Mice were injected 0.1 mM/kg of gadolinium-diethylene-triamine-pentaacetic acid (Gd-DTPA, MW= 938 Da; Bayer Healthcare) as a bolus injection in the tail vein, followed by imaging. DCE-MRI was performed using a transverse fast T1 mapping that consisted of obtaining pre-contrast (three sequences) and post-contrast (15 sequences) images up to 15 min after the contrast injection. The details of the pulse sequence T1 RARE for T1 weighted imaging are: FOV = 2 cm × 2 cm, slice thickness = 1.5 mm, slice gap = 0, matrix size = 128 × 128, TR/TE =377 ms/12.3 ms, number of averaging = 2, total scan time = 48 s 25 ms.

#### MRI Data Analysis and Processing

The method is based on the leakage of the contrast agent from plasma compartment into brain compartment through BBB resulting in a change of the MRI signal intensity. The rate of changes in the MRI signal intensity relates to the BBB permeability (*Ki*). The T1 map was reconstructed with the t1epia fitting function in the Bruker ParaVision Image Sequence Analysis (ISA) tool. Previous research has demonstrated that the blood-to-tissue transfer or influx constant, *Ki*, could be obtained by a graphical analysis of time series of tissue and arterial contrast agent concentration.^57^ Since the contrast agent concentration is proportional to changes of 1/T1(Δ(1/T1(t))), the color-coded map of *Ki* was constructed from repeated estimates of Δ(1/T1(t)), where pixels with higher intensity color represent higher BBB permeability. A custom-made computer program in MATLAB (Mathworks, Massachusetts, USA), which implemented the above principle, was used to generate the *Ki* map.

### The assessment of the BBB permeability to GM_1_-liposomes

Fluorescently-labeled GM_1_-liposomes were constructed on the basis of a matrix of egg yolk phosphatidylcholine (Lipoid GmbH, Germany) with a content of 10 mol. percent of ganglioside GM_1_ from bovine brain. Purified ganglioside GM_1_ from bovine brain was kindly provided by Dr. Ilya Mikhalyov (Shemyakin-Ovchinnikov Institute of Bioorganic Chemistry RAS). GM_1_-liposomes were produced by a standard extrusion method after hydration of the lipid film.^58^ Liposomes contained 1 mol.% BODIPY-phosphatidylcholine in the bilayer (the structure is shown in Figure 3 in SI). A high-efficient fluorescent probe (*λ*_ex_ = 497 nm, *λ*_em_ = 504 nm), synthesized as described earlier,^59^ was kindly provided by Dr. Ivan Boldyrev (Shemyakin-Ovchinnikov Institute of Bioorganic Chemistry RAS). After extrusion through membrane filters with a pore diameter of 100 nm (Extruder Lipex, Northern Lipids, Canada) liposomal dispersions in physiological saline (phosphate buffer, pH 7.1, total lipid concentration 25 mM) were produced. Measured by dynamic light scattering technique (BI9000, Brookhaven Instruments, USA), the average diameter and polydispersity index for liposomes were 104 nm and 0.076, respectively.

#### Immunolabeling

The brains were fixed with 4% neutral buffered formalin. After fixation, the brains were cryoprotected using 20% sucrose in phosphate buffered saline (PBS) (10 ml/brain mouse) for 48h at 4°C. The brains were frozen in hexane cooled to –32 ÷ –36°C. Crysection (14 μm) of parietal cortex were collected on poly-L-Lys, Polysine Slides (Menzel-Glaser, Germany) using cryotome (Thermo Scientific Microm HM 525, Germany) and a liquid for fixing a Tissue-Tek sample (Sakura Finetek, USA). Cryosections were blocked in 150 μl 10% BSA/PBS for 1h, then incubated overnight at 4°C with 130 μl anti-rat SMI-71 mouse antibody (1:500; Santa Cruz Biotechnology, Santa Cruz, CA, USA); anti-rat GFAP rabbit antibody (1:500; Sigma-Aldrich Corporation, St Louis, CIIIA). After several rinses in PBS, the slides were incubated for 1h with 130 μl fluorescent-labeled secondary antibodies on 1% BSA/PBS (1:500; Santa Cruz Biotechnology, Santa Cruz, CA, USA, Alexa Four 555; Invitrogen, Molecular Probes, Eugene, OR, USA). The confocal microscopy was performed using the microscope Nikon TE 2000 Eclipse, Tokyo, Japan.

The GM_1_-liposomes in physiological saline were injected to mice (0.2 ml/100 g, i.v.) in following groups: 1) control, no sound (n=10); 2) in 1h (n=10), in 4h (n=10) and in 24h (n=10) after the sound exposure.

### Biochemical assays of plasma epinephrine

The plasma epinephrine level (ng/ml) was determined using ELISA kits (Abnova, Taiwan) at normal state (before sound), during sound stress (at the last minute (120 min) of sound stress) and in post-stress-period (1h after sound exposure) in mice (n=10 in each group). The plates were read at 450 nm using an ELx 800 plate reader (BioTek Instruments Inc.). The detection limits were 0.3 ng/ml (with intra and inter assay variation coefficients of 11.2-16.3% and 8.7-12.6%, respectively).

### Laser speckle contrast imaging of the cerebral blood flow (CBF)

A custom-made laser speckle contrast imaging system was used to monitor the effects of adrenaline on the CBF at normal state and in 1h/24h after sound exposure in 7 mice. Light from a single mode He-Ne laser (Thorlabs, HNL210L, 632.8 nm, 21 mW) was launched into the single mode polarization maintained optical fiber (Thorlabs, P3-630PM-FC-1) in order to achieve a flexible and uniform illumination of the sample. The angle of incidence was set to approx. 45 deg. The resulting fluctuating speckle pattern was imaged using ComputarM1614-MP2 lens with one extension ring in order to achieve near unity magnification. F-stop for the lens was set to f/6 to get the speckle-to-pixel ratio about 15. Complementary a metal-oxide-semiconductor (CMOS)-camera Basler acA2500-14 gm with pixel area of 2.2 x 2.2 μm^2^ was used as a detector. The frame rate of the recorded sequence was 40 frames per second. The spatial speckle contrast was calculated using the ratio between the standard deviation and the mean value of the intensity fluctuations within 5×5 pixels sliding windows. Fifty consecutive speckle contrast frames were averaged in order to improve the signal-to-noise ratio. The automated segmentation algorithm, described in our previous work,^60^ was used to calculate the mean value of CBF at macro- and micro-levels.

### Assessment of cell blebbing

Rat brain cortex cells were obtained accordingly to the protocol of mechanical dissociation in phosphate buffered saline (PBS) (10 ml/rat brain).^61^ The resulting cell suspension (100 μl) was exposed to the sound (5-10-20-40 min) or was left at room temperature for the same period as a control. Immediately after exposure, 10 μl of the cell suspension were transferred to the slide and covered with a cover slip. The analysis of blebbing was carried out accordingly to the protocol of Inzhutova et al.^62^ A phase contrast microscopy (40 X microscope objective) was performed with an Olympus BX-41 microscope, the images were taken with an Olympus DP72 camera. The parameters such as the number of intact cells, the number of cells in the state of initial or terminal blebbing, and the number of necrotic cells were evaluated. At least 10 fields of view were analyzed and the counting was performed in 100 cells.

### Measurement of transendothelial electrical resistance (TEER)

The TEER was measured using an EVOM2 volt-ohmmeter with the STX-2 chopstick electrodes (World Precision Instruments, Sarasota, FL, USA). Resistance values (Ωcm^2^) were corrected by subtracting the resistance of an empty Transwell filter. At each time point (0-0.5-1.5-4-24 h) TEER measurements were taken.

The experiment was performed using *in vitro* model of BBB. The exposure time was 5 min (Sound time 5 min, n=6) and 10 min (Sound time 10 min, n=6). The control group was not exposed to sound (no sound, n=6).

### Establishment of BBB model *in vitro*

Brain microvessel endothelial cells (BMECs) were isolated from the brain of Wistar rats (postnatal day 10-14). Isolation and establishment of primary culture of BMECs were done according to the protocol of Liu et al.^63^ The BMECs obtained were phenotyped by standard immunohistochemistry protocol using primary anti-ZO1 antibodies (Santa Cruz Biotechnology, Inc., sc-8147) and secondary antibodies labeled with Alexa Fluor 488 (Abcam, ab150117) followed by the detection with the fluorescent microscope. Astroglial and neuronal cells for the BBB mode were obtained from embryonic rat neurospheres as described in^64^. The integrity of the in vitro BBB model containing BMECs monolayer grown on inserts, astrocytes and neurons grown in the well was confirmed with TEER measurements.

### Immunohistochemical assay

Animals (Wistar rats, 270-300 g) in the control group (before sound, n=7) and experimental groups in 1h and 24h after sound (n=7 in each group) were sedated (pentobarbital 50 mg/kg, intraperitoneal), intracardially perfused with PBS followed by 4% paraformaldehyde, and brains were removed. Brain slices (50 μm) were processed according to the standard protocol for immunohistochemistry with the appropriate primary and secondary antibodies and visualization with confocal laser microscopy Olympus FV10i-W (Olympus, Japan).

The expression of antigens on free-floating sections was evaluated using the standard method of simultaneous combined staining (Abcam Protocol). Brain slices were blocked in 150 μl 10% BSA/0.2% Triton X-100/PBS for 2 h, then incubated overnight at 4°C with Rb antimouse Anti-beta 2 Adrenergic Receptor antibody (1:500; Abcam, ab182136, Cambridge, USA); Rb anti-mouse Anti-beta Arrestin 1 antibody (1:500; Abcam, ab32099, Cambridge, USA); CLND-5 (1:500; Santa Cruz Biotechnology, sc-28670, Santa Cruz, USA); ZO-1 (1:500; Santa Cruz Biotechnology, sc-8147, Santa Cruz, USA); MMP9 (1:500; Abcam, ab58803, Cambridge, USA); HIF-1 (1:500; Abcam, ab2771, Cambridge, USA); VEGF (1:500, Abcam, ab1316, Cambridge, USA) followed by 2 h at room temperature. After several rinses in PBS, the slides were incubated for 3h at room temperature with fluorescent-labeled secondary antibodies on 1% BSA/0.2% Triton X-100 /PBS (1:500; Goat A/Rb, Alexa 555-Abcam, UK, ab150078). Confocal microscopy of the mice cerebral cortex was performed using a fully automated confocal laser scanning microscope with water immersion Olympus FV10i-W (Olympus, Japan). ImageJ was used for image data processing and analysis. The areas of expression of antigens were calculated using plugin “Analyze Particles” in the “Analyze” tab, which calculates the total area of antigen-expressing tissue elements - the indicator “Total Area”. In all cases, 10 regions of interest were analyzed.

### Assessment of apoptosis with TUNEL method

The number of apoptotic cells was evaluated with the TUNEL method using the “17-141 TUNEL Apoptosis Detection Kit” (Abcam, UK) in accordance with the standard protocol provided by the manufacturer. The brain slices were incubated in a reaction mixture with TdT and nucleotides labeled with avidin-FITC. To stain the nuclei of cells, propidium iodide with RNase from another set was added for 15 min at room temperature. Confocal microscopy was performed with an Olympus FLUOVIEW FV10i-W (Tokyo, Japan). An Olympus FLUOVIEW Viewer 4.0 (Olympus, Japan) or ImageJ (free software) were used for images processing and analysis. FITC-positive (apoptotic) cells were counted as a percentage of the total number of cells. Analysis of apoptosis with TUNEL protocol in the brain slices was performed in 1h after sound exposure during 2.5, 5, 10, 20 min, n=7 in each group (See Figure 6 in SI).

For **histological analysis of brain tissues and vessels**, four groups of animals were used: 1) control, no sound; 2) at 1h, 4h and 24h after sound exposure, respectively, n=15 in each group. All mice were decapitated after the performed experiments. The samples were fixed in 10% buffered neutral formalin. The formalin fixed specimens were embedded in paraffin, sectioned (4 μm) and stained with hematoxylin and eosin. The histological sections were evaluated by light microscopy using the digital image analysis system Mikrovizor medical μVizo-103(LOMO, Russia).

### The assessment of hearing loss and cognitive impairment (Fear conditioning test)

To eliminate nonspecific influence (hearing loss, cognitive impairment) of sound (100 dB, 370 Hz) we used *Fear conditioning* (FC) test.^27^ FC is widely used as a test for memory consolidation and is based on auditory and contextual stimuli^31^. The protocol included three stages: 1) training, 480 sec (three times of stimulation with 30 sec (so-called “white noise”, 55 dB) and a 0.1-0.8 mA footshock during the last 2 sec of the sound); 2) context test, 300 sec (without any stimulation); 3) cued test, another testing chamber with very different properties, providing a new context, 360 sec (with “white noise”, 55 dB, for last 180 sec). Additional stimulation with the sound (100 dB) for 1 min was carried out at 1 or 24 h before the third stage initiation in the experimental group. N=5 mice in each group. FC is widely used as a test for memory consolidation and is based on auditory and contextual stimuli.^28^ Freezing response which is evaluated in the FC depends on memory quality, sound recognition/hearing deficit, and may also reflect some neurobehavioral changes in animals, i.e. anxiety.^29^

### Statistical analysis

The results are presented as mean ± standard error of the mean (SEM). Differences from the initial level in the same group were evaluated by the Wilcoxon test. Intergroup differences were evaluated using the Mann-Whitney test and ANOVA-2 (post hoc analysis with Duncan’s rank test). The significance levels were set at p < 0.05-0.001 for all analyses.

## Acknowledgments

O.S.-G., D.B., A.S., N.N., A.A., M.U., J.K. were supported by Grant of Russian Science Foundation *№* 17-15-01263. D.B. was supported by NIH R21NS091600 and P20GM109089. D.Z., C.Z., W.F. were supported by National Natural Science Foundation of China (NSFC) (Grant Nos. 31571002, 81701354), the Foundation for Innovative Research Groups of the National Natural Science Foundation of China (Grant No. 61721092), and Director Fund of WNLO. V.T. and O.S.-G. were supported by grants RFBR *№* 17-02-00358 and MES RF *№* 17.1223.2017/AP.

## Author contributions

O.S.-G., V.Ch., D.Z., Q.L., V.T. and J.K. conceived of the presented idea and planned the experiments. D.B., O.B., E.V., A.A., V.S., A.M., N.M., E.O., E.B., A.T., A.S., N.N., Y.Y., C.Z, W.F., A.A., M.U., N.S., A.K., A.T., A.E.S. and A.P. carried out the experiments. O.S-G., J.K., and V.T. contributed to the interpretation of the results. O.S.-G. took the lead in writing the manuscript. All authors discussed the results and contributed to the final manuscript.

