## Supplementary material for "Loud music and the specific sound stress open the blood-brain barrier: new fundamental, biomedical, and social aspects": Tables and figures

*Corresponding author

**Table 1 - The effect of loud music and sound on BBB permeability to the EBD-albumin complex (µg/g tissue)**

| **Sound level (dB) and time elapsed after sound exposure (h)** | **Content of EBD (µg/g tissue)** | |
|---|---|---|
| No sound (the control group) | 0.15±0.01 | |
| | **Music** | **Sound** |
| 100 dB | | |
| 1 h | **2.60±0.06** *** | **2.80±0.08** *** |
| 4 h | 0.19±0.03 | 0.17±0.07 |
| 24 h | 0.16±0.03 | 0.14±0.02 |
| 90 dB | | |
| 1 h | **2.70±0.04** *** | **2.20±0.09** *** |
| 4 h | 0.15±0.03 | 0.18±0.07 |
| 24 h | 0.19±0.07 | 0.16±0.09 |
| 70 dB | | |
| 1 h | 0.17±0.08 | 0.16±0.06 |
| 4 h | 0.19±0.06 | 0.18±0.04 |
| 24 h | 0.19±0.09 | 0.15±0.08 |

$p<0.001$: *** - vs. before sound (the control group), n=15 in each group.

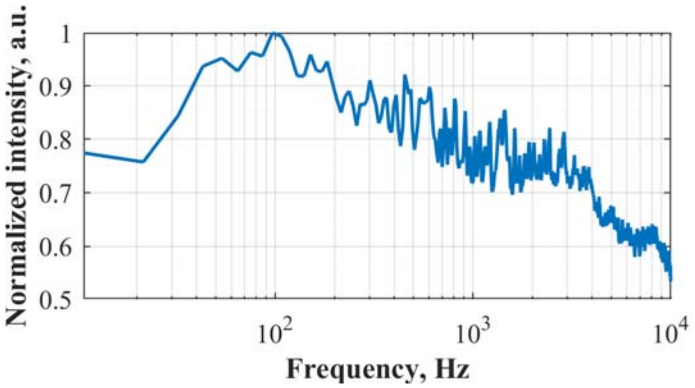


**Figure 1 – The frequency range of music (Scorpions Still Loving You")**: frequencies in the range of 11-10,000 Hz and maximal intensity around 100 Hz.

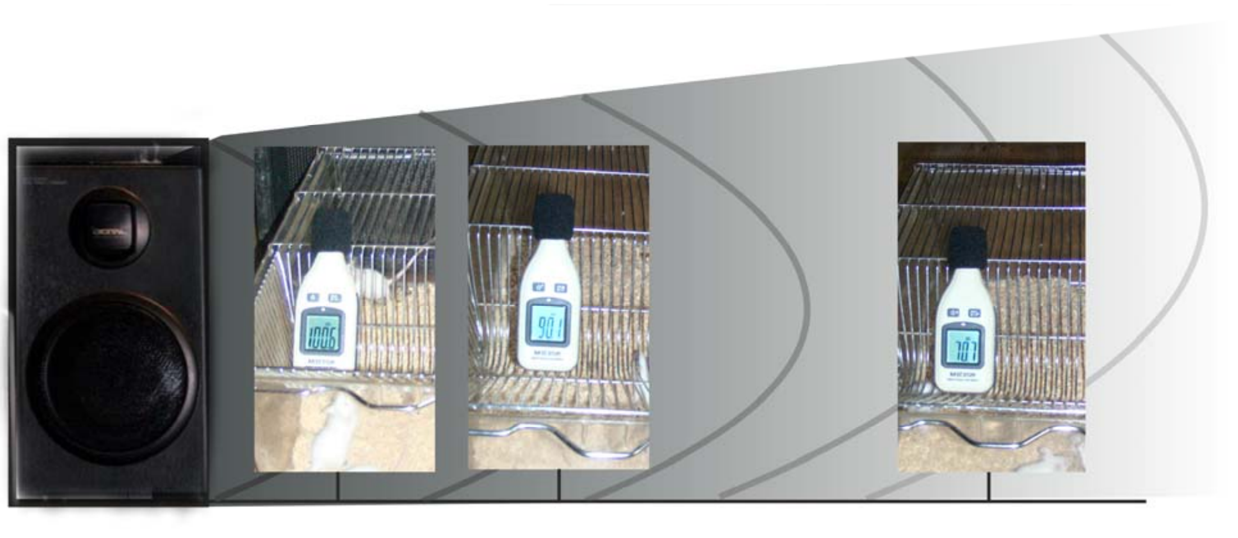

**Figure 2 – Design of experiment for the study of effects of loud music (100 dB, 11-10,000 Hz, Scorpions "Still loving you") on BBB.** To produce the loud music, we used loudspeaker generated sound 100 dB. The sound energy was measured directly in cage of animals using sound level meter. The same design of experiments was for the study of loud sound (100 dB, 370 Hz).

A

B

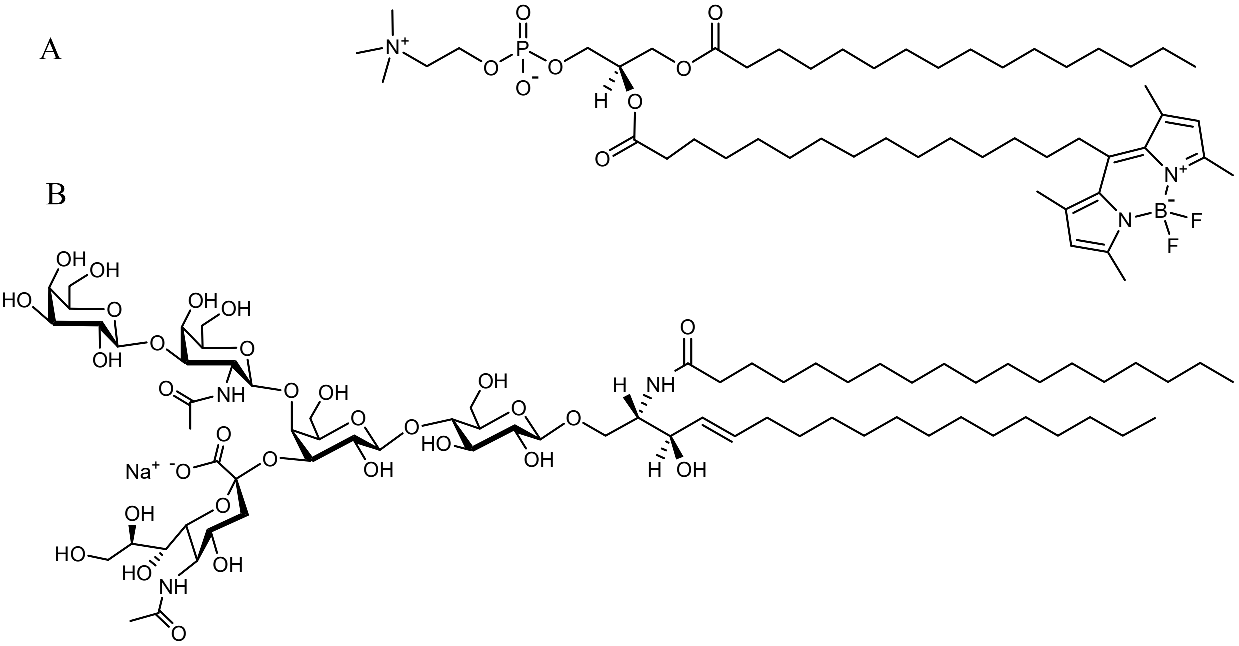

**Figure 3 - Molecular structures of lipid bilayer components of fluorescently labelled $GM_1$-liposomes based on egg phosphatidylcholine**: A - BODIPY-phosphatidylcholine {1-palmitoyl-2-[15-(4,4-difluoro-1,3,5,7-tetramethyl-4-bora-3*a*,4*a*-diaza-*s*-indacene-8-yl)pentadecanoyl]-*sn*-glycero-3-phosphocholine}; B - representative structure of ganglioside $GM_1$ from bovine brain.

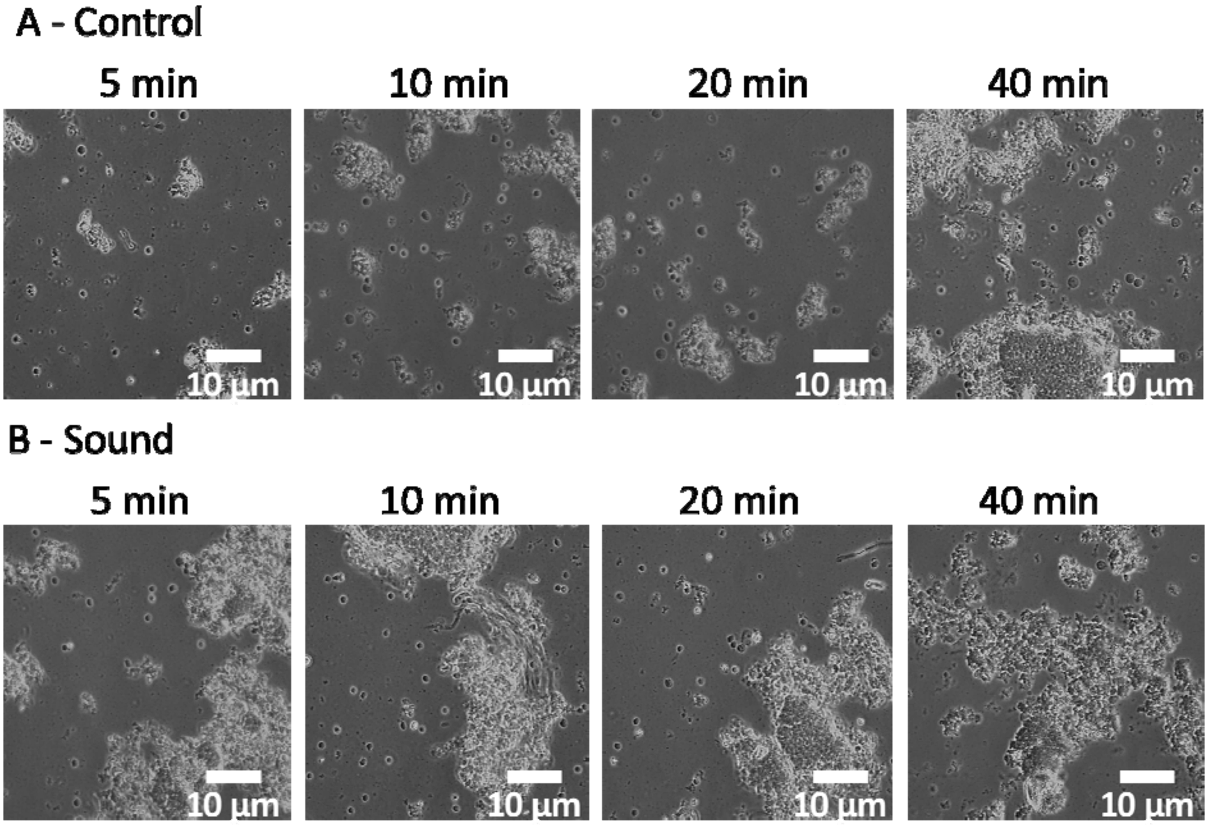


**Figure 4 – The study of appearance of blebbing brain cells exposed to sound during 5, 10, 20, 40 min.** The signs of initial blebbing: presence of small vesicles on the cell membrane (up to 1/3 of the cell radius); the signs of terminal blebbing: presence of large membrane blisters (more than 1/3 of the cell radius). 5-10 min exposure to sound resulted in cell membrane blebbing, whereas longer exposure (up to 20 min) resulted in significant cell destruction. Bars represent 10 µm. At least 10 fields of vision were analyzed and the counting was carried out in 100 cells.

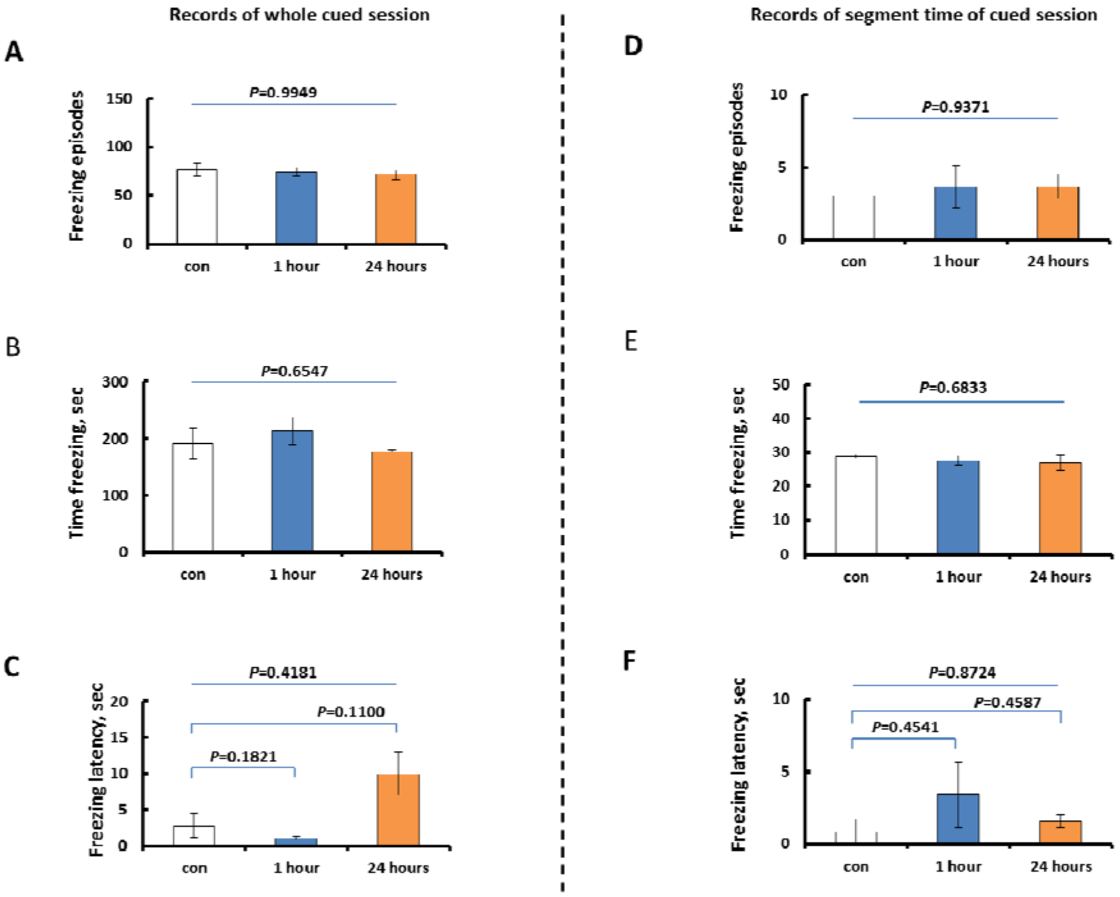


**Figure 5. Cognitive function and hearing ability after additional high tone stimulation in the cued stage of *Fear Conditioning* test.** Total freezing episodes (**A**), time freezing (**B**) and freezing latency (**C**) during whole cued session (360 s, left panel) are presented as a total score of cognitive function and hearing ability. Freezing episodes (**E**), time freezing (**F**) and freezing latency (**D**) during selected time segment (duration – 30 s, start and point - 180 and 210 s of time record, respectively) are presented as a score of hearing ability according to "white noise" appearance ("white noise" starts from 180 s of time record, right panel). **A**) Total freezing episodes, one-way ANOVA, $P$=0.9949, $F_{2,15}$= 0.0051; **B**) total time freezing, one-way ANOVA, $P$=0.6547, $F_{2,15}$= 0.4376; **C**) freezing latency, one-way ANOVA ($P$=0.4181, $F_{2,15}$=0.9331) with following *Bonferroni post-hoc* test; **D**) freezing episodes in selected time segment, one-way ANOVA, $P$=0.9371, $F_{2,15}$=0.0657; **E**) time freezing in selected time segment, one-way ANOVA, $P$=0.6833 , $F_{2,15}$=0.3921; **F**) Freezing latency in selected time segment, one-way ANOVA ($P$=0.8724, $F_{2,15}$=0.1374) with following *Bonferroni post-hoc* test. N = 5 mice in each group. Con – control group, 1 hour and 24 hours – experimental groups with additional stimulation with high frequency sound for 1 min (100dB) at 1 or 24 hours before the cued stage initiation.

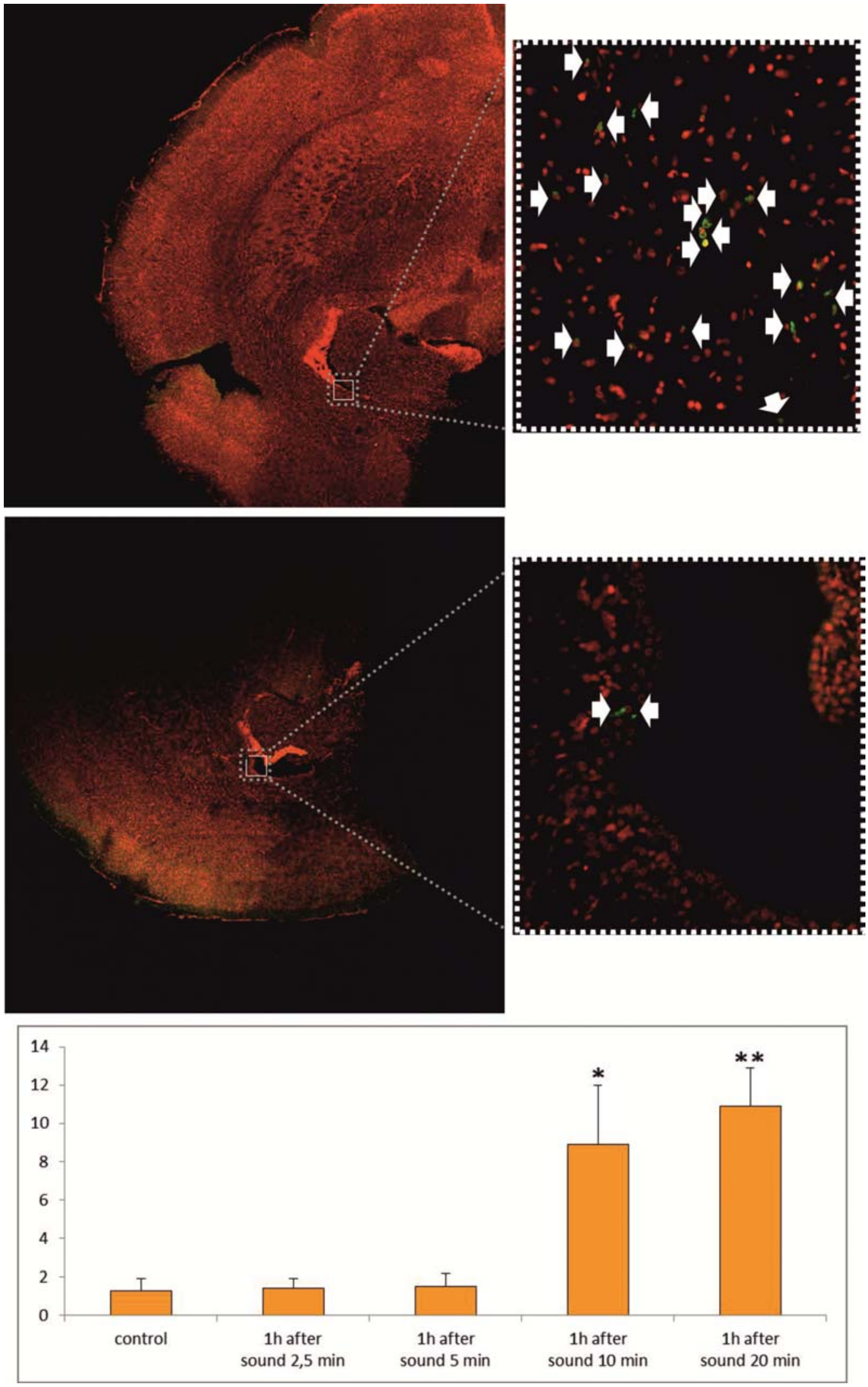


**Figure 6 – Assessment of apoptosis with TUNEL protocol in brain slices *in vitro*.** 2.5 and 5 min exposure to sound does not induce significant changes in the appearance of apoptotic cells in the brain cortex. 10 min and 20 min sound exposure results in significant increase in TUNEL-positive cells in 1 h after the sound action as shown on photos (lower panel – control, upper panel – in 1 h after 10 min sound exposure, arrows indicate TUNEL-positive cells). * - $p<0.05$; ** - $p<0.01$ vs. control group (no sound action), n=7 in each group.

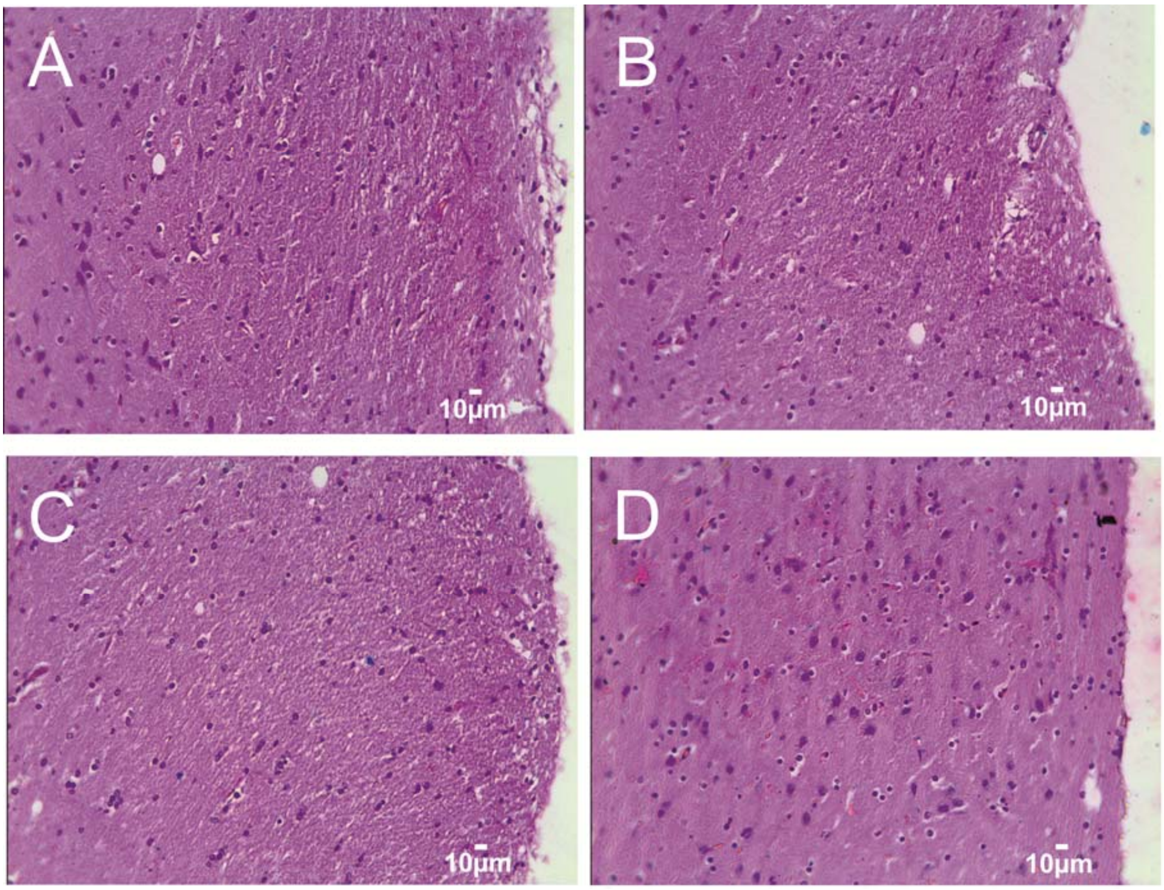


**Figure 7 - Histological analysis of brain tissues before and after sound exposure**: A – the control group, no sound influences, B, C and D – in 1h, 4h and 24h after sound exposure, respectively. No changes in the brain tissues and vessels were found in 1h, 4h and 24h after sound impact when BBB was opened and closed, respectively (n=15 in each group). Hematoxylin & Eosin staining. Bars represent 10 µm (246.4X).
